# Anti-biofilm Activity of Graphene Quantum Dots via Self-Assembly with Bacterial Amyloid Proteins

**DOI:** 10.1101/550285

**Authors:** Yichun Wang, Usha Kadiyala, Zhibei Qu, Paolo Elvati, Christopher Altheim, Nicholas A. Kotov, Angela Violi, J. Scott VanEpps

## Abstract

Bacterial biofilms represent an essential part of Earth’s ecosystem that can cause multiple ecological, technological and health problems. The environmental resilience and sophisticated organization of biofilms are enabled by the extracellular matrix that creates a protective network of biomolecules around the bacterial community. Current anti-biofilm agents can interfere with extracellular matrix production but, being based on small molecules, are degraded by bacteria and rapidly diffuse away from biofilms. Both factors severely reduce their efficacy, while their toxicity to higher organisms create additional barriers to their practicality. In this paper we report on the ability of graphene quantum dots to effectively disperse mature *Staphylococcus aureus* biofilms, interfering with the self-assembly of amyloid fibers - a key structural component of the extracellular matrix. Mimicking peptide-binding biomolecules, graphene quantum dots form supramolecular complexes with phenol soluble modulins, the peptide monomers of amyloid fibers. Experimental and computational results show that graphene quantum dots efficiently dock near the *N*-terminus of the peptide and change the secondary structure of phenol soluble modulins, which disrupts their fibrillation and represents a novel strategy for mitigation of bacterial communities.

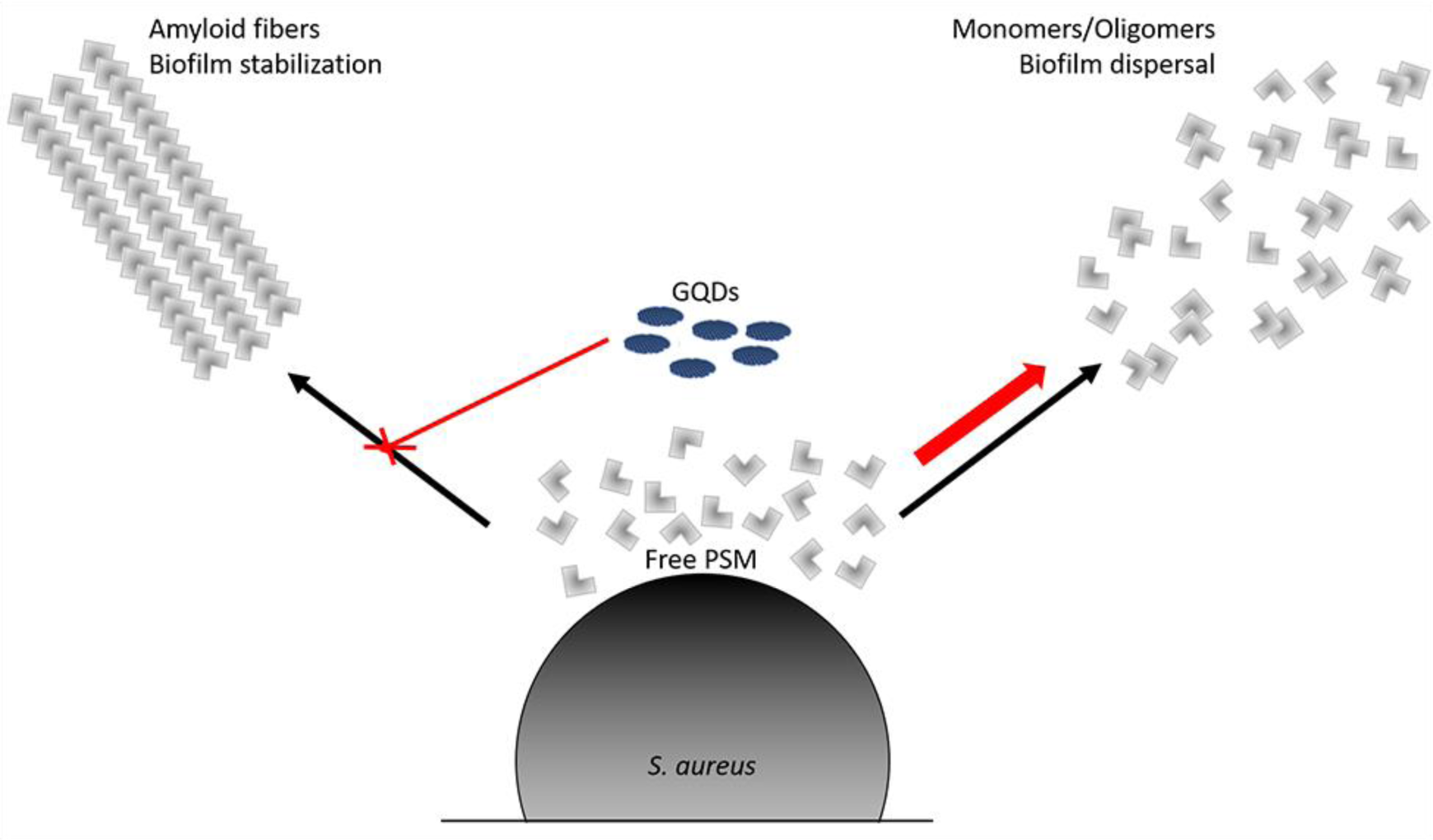
GQD mediated staphylococcal biofilm dispersal. GQDs interact with PSM peptides and frustrate the fibrillation process. The reduction in amyloid fibers prevents robust stabilization of the biofilm. In addition, there is an increase in free monomeric and oligomeric PSM peptides which trigger dispersal events.

---

Most microorganisms survive in the form of biofilms - structurally and functionally sophisticated microbial communities. Biofilms are implicated in numerous technological problems from water membrane failures to airplane crashes^1,2^ as well as in various diseases from persistent infections to cancer^3,4^. Biofilms also form a large component of the ecosphere on Earth and their dysregulation can also cause significant ecological consequences.^5^

Biofilms act like multicellular organisms and possess exceptional adaptability and resilience to environmental factors and pharmaceuticals.^6,7^ Both of these properties are afforded by the protective function of extracellular matrix (ECM), which is made from several types of biomolecules (*i.e.*, polysaccharides, DNA, and peptides).^8,9^ To date only a few classes of compounds have been shown to be effective anti-biofilm agents.^10,11^ Peptides rich in amino acids such as proline, arginine, phenylalanine or tryptophan have shown broad-spectrum activity in killing bacteria within biofilms by targeting a universal stringent stress response in bacteria.^12^ However, they are currently being developed only as antibiotic adjuvants because they are subject to rapid proteolytic degradation and are relatively expensive to synthesize. Small molecules have been used to inhibit the quorum sensing (QS) system^13^ or interfere with the polymerization of amyloid proteins.^14^ While further exploration of these strategies of biofilm inhibition need to be explored, the rapid loss of the small molecules due to diffusion into environment reduce their efficacy and increase their environmental toxicity.

These problems with the design of effective anti-biofilm agents prompted us to search for new materials platforms. Graphene quantum dots (GQDs) are a single layer graphene a few nanometers in diameter. They are often regarded as biocompatible alternatives of II-VI semiconductor NPs also known as quantum dots (QDs).^15^ GQDs have been investigated for antibacterial activity due to reactive oxygen species production and membrane disruption.^16–18^ While these functionalities highlight the significance of GQDs as a part of microbiology toolbox, we hypothesize that these nanoscale particles can interfere with the self-assembly processes of biomolecular components of the biofilms. Representing the common tendency of nanoscale particles to self-assemble, this hypothesis is grounded in the multiple observations of GQDs and other NPs to specifically interact and assemble with other nanoscale particles and biomolecules.^19–27^ As applied to biofilms, GQDs can form ‘decoy’ complexes with the key structural components of ECM, thereby inhibiting the biofilm formation. Recently this functionality was confirmed by observation of GQDs acting as inhibitors of fibrillation of the protein characteristic for Alzheimer’s and Parkinson’s disease.^28–31^ Counteracting known problems of other anti-biofilm agents mentioned above, GQDs have high molecular weight (10^3^ to 10^5^ g/mol) that slows diffusion and are resistant to proteolytic degradation.

As an experimental model to study the anti-biofilm effects of GQDs, we used biofilms of *Staphylococcus aureus* -- a major cause of hospital- and community-associated bacterial infections in the U.S. and around the world.^32^ Formation of *S. aureus* biofilms on host tissues and implanted medical devices contributes to chronic infections, as biofilms are known to be exceptionally resistant to host immune response and tolerant to antibiotics.^36^ The formation and dispersion of staphylococcal biofilms is dependent on the secretion of the phenol-soluble modulins (PSMs)^32–34^--small α-helical amphipathic peptides that have previously been implicated in bacterial virulence.^35,36^ Assembly of PSMs into fibers^35^ promotes the maturation of ECM,^32,37,38^ whose integrity renders these bacterial communities resistant to dispersal by proteinase K, Dispersin B, DNase and sodium dodecyl sulfate.^32^

The self-assembly process of PSMs into amyloid-like fibers involves backbone hydrogen bonding and side-chain interaction (*e.g.*, hydrophobic interaction, π-stacking, and van der Waals attraction).^39^ We took advantage of the characteristics of GQDs including polarizability, amphiphilic character, ability to form hydrogen bonds and participation in π-π stacking.^40^ Based on microscopy, spectroscopy, and computational data, we found that the growth of fibers from PSM peptides was dramatically stunted in the presence of GQDs, which resulted in dispersal of mature *S. aureus* biofilms. Computer simulations indicate that GQD docks at the *N*-terminal of PSM peptides via carboxyl edge-group and alters the PSM conformation.

## RESULTS AND DISCUSSION

GQDs used in this study had a diameter of 2-8 nm^15^ and carried nine carboxyl edge-groups on average. (**Figure S1**) To investigate how GQDs affect amyloid-rich biofilms, *S. aureus* biofilms were grown in peptone-NaCl-glucose (PNG) media for four days in flow cells.^32^ *S. aureus* biofilms were also grown in tryptic soy broth supplemented with glucose (TSBG) media that inhibits amyloid synthesis as a negative control. Confocal images showed that *S. aureus* formed biofilm covering the entire surface of the flow cell in both TSBG and PNG without GQDs (**Figure 1a, d**). In the presence of 50-μg/mL GQDs, the biofilm grown on PNG was largely dispersed whereas the biofilm on TSBG retained its structure (**Figure 1b, e**). A further dispersion of biofilm structure occurred in the presence of 500-μg/mL GQDs in PNG condition, while TSBG biofilm remaining fully present (**Figure 1c**, **f**). According to quantitative image analysis, porosity of the PNG biofilm (**Figure 1g**) increased from 29.4 ± 8.8% to 50.7 ± 6.9% when cultured with 50-μg/mL GQDs; the porosity further increased to 63.4 ± 5.0% for 500-μg/mL GQDs. The thickness of the PNG biofilm **(Figure 1h)** decreased from 21.9 ± 2.5 μm to 16.9 ± 3.4 μm when cultured with 50-μg/mL GQDs; the thickness further decreased to 12.7 ± 3.4 μm with 500-μg/mL GQDs. The thickness for biofilms grown in TSBG media showed limited change with addition of any dose of GQDs **(Figure S2)**. The structural changes of amyloid-rich biofilms before and after GQD (500 µg/mL) treatment was further verified by scanning electron microscopy (SEM) (**Figure 1i, j**). Here the change in porosity of the PNG biofilm is even more distinct than that in confocal microscopy, which increased to 71.2 ± 6.6 % (**Figure 1k**). This is likely a result of dehydration process required for SEM sample preparation, which washed off weakly attached bacterial cells and loose ECM content. The phenomenon not only confirmed the increase of biofilm porosity after GQD treatment but also indicated reduced integrity of remaining biomass.

**Figure 1.**
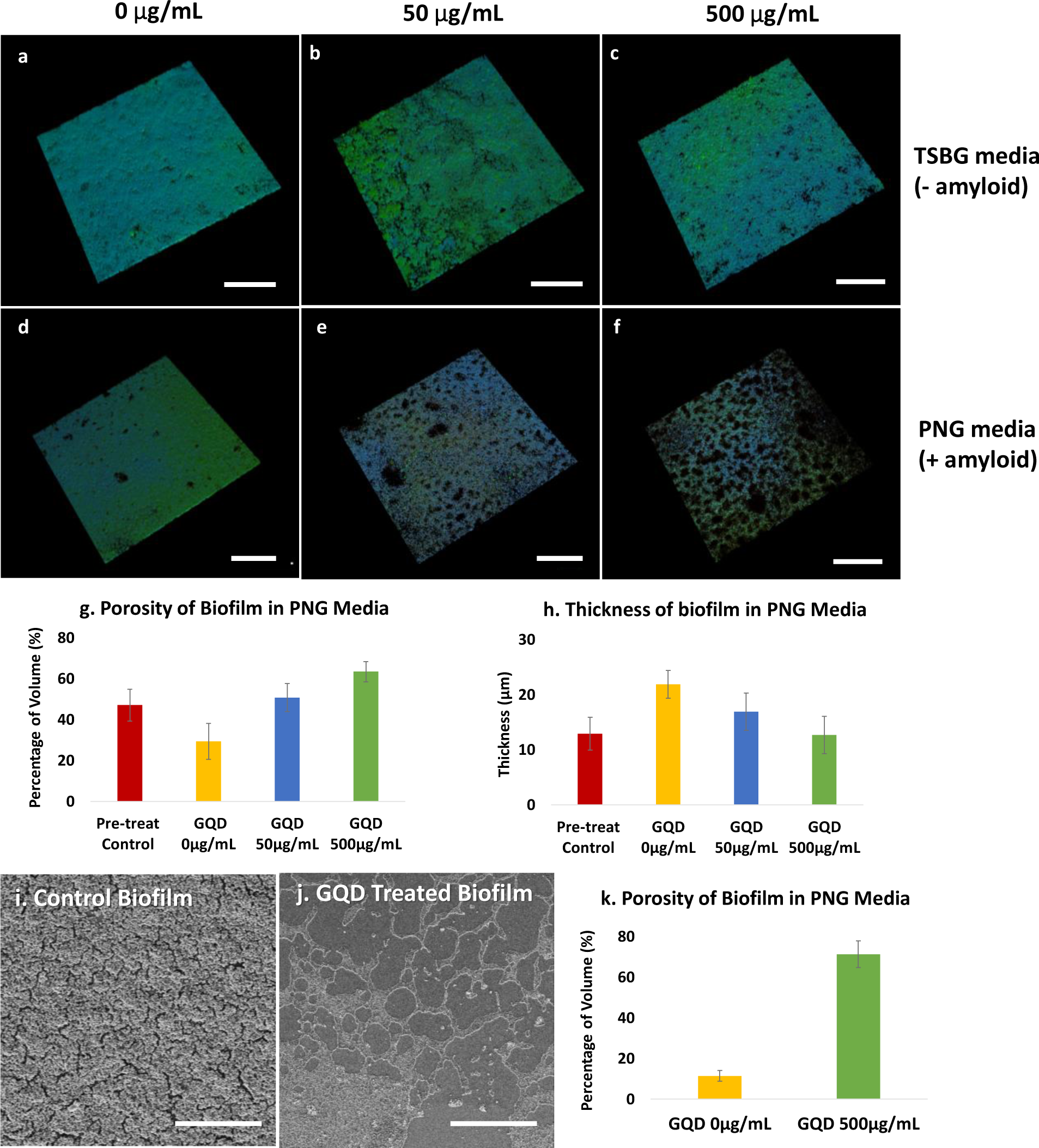
Effects of GQDs on *S. aureus* biofilms. **(a-f)** Confocal microscopy of *S. aureus* biofilm grown for 3 days and then treated for 1 day; stains - polysaccharide intercellular adhesin (PIA; green) and bacterial cells (blue). Biofilms grown in TSBG medium in presence of **(a)** 0 µg/mL **(f)** 50 µg/mL and **(c)** 500 µg/mL GQDs. Biofilms grown in PNG medium in presence of **(d)** 0 µg/mL **(e)** 50 µg/mL and **(f)** 500 µg/mL GQDs. Bar graphs of quantified **(g)** porosity and **(h)** thickness of biofilms analyzed from the *z*-stack images for each group in **(a-f)**. **(i, j)** Representative SEM images and their **(k)** calculated porosity of *S. aureus* biofilms grown in PNG. Scale bar: 200 μm **(a-f)**, 50 μm **(i, j)**.

Of note, the growth rate of planktonic *S. aureus* did not differ significantly with or without GQDs up to 200 μg/mL (**Figure S3**). This observation indicates that the GQDs are not toxic to the individual bacteria cells *per se*, and thus the biofilm dispersal is associated with change in ECM integrity only. Furthermore, the specificity for the GQD effect on PNG grown biofilm versus the TSBG biofilm suggests that amyloid is the unique target of GQDs among other components of the ECM.

The formation of amyloid fibers in *S. aureus* biofilms grown in PNG were confirmed by transmission electron microscopy (TEM). The fibers, with a diameter of about 12 nm, were closely associated with bacterial cells and surrounding ECM; their dimensions are comparable to those observed in prior studies^32^ (**Figure 2a**). ECM was extracted from the biofilm cultures and further analyzed with respect to the effect of GQDs on ECM morphology. In general, various components from ECM may closely combine with the amyloid-fibril core (**Figure 2a, b**), which may contain PSM peptides, extracellular DNA and polysaccharide intercellular adhesion (PIA). The GQDs, with concentration of 50 µg/mL, dramatically altered the morphology and integrity (**Figure 2c**).

**Figure 2.**
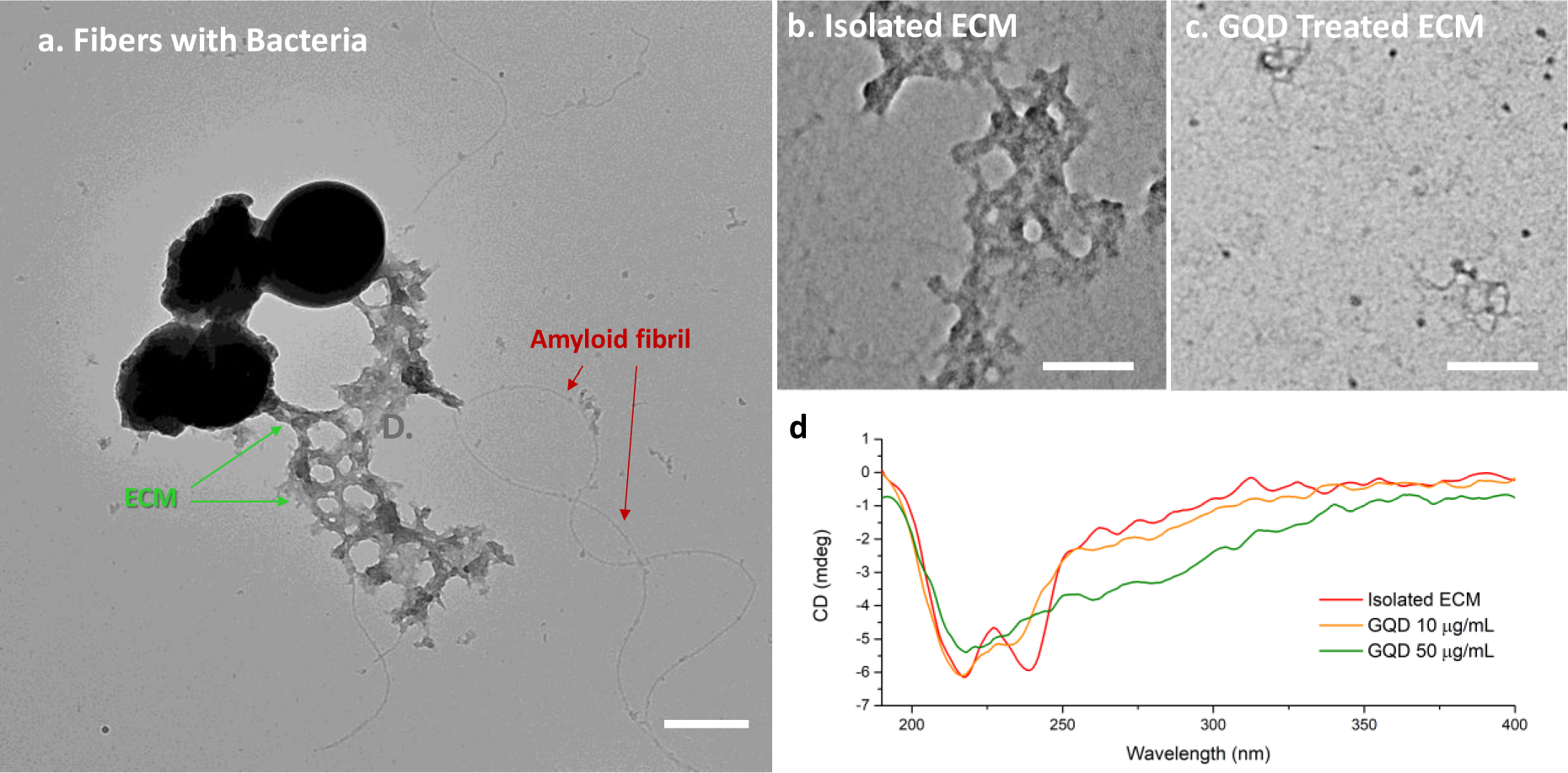
TEM and CD spectra of isolated ECM from *S. aureus* biofilm grown in PNG and exposed to GQDs. **(a)** Amyloid fibrils in composition with ECM attached to bacteria; **(b)** Extracted composite ECM treated with **(b)** 0 µg/mL and **(c)** 50 µg/mL GQDs for 4 days. **(d)** CD spectra of isolated ECM treated with 0 µg/mL, 10 µg/mL, and 50 µg/mL GQD for 4 days. Scale bars: 500 nm **(a)**, 100 nm **(b, c)**. Only low concentrations of GQDs were evaluated in this test due to the low concentration of ECM obtained from the isolation procedure compared to that from intact biofilms.

Circular dichroism (CD) spectrometry was used to obtain complementary information about the secondary structure of the amyloid fibrils. A wide negative peak between 200 nm to 230 nm observed in the CD spectrum (**Figure 2d**) is characteristic of amyloid-like fibrils including typical β-sheet signals at negative at 218 nm and α-helix negative bands at 208 nm and 222 nm. The negative CD peak at ≈245 nm is associated with extracellular DNA.^41^ All peaks between 200 nm to 250 nm were significantly decreased after two-hour incubation with GQDs suggesting that GQDs cause disruption of the fibril composite structure of extracted biofilm ECM within this timeframe.

The main content of the isolated amyloid fibers from PNG biofilm consists of small α-helical amphipathic peptides, PSMs.^32^ PSMα1 is one of these PSM peptides with capacity to form amyloid-like fibrils. We used commercially available, synthetic PSMα1 to further investigate the effects of GQDs. Their fibrillation was monitored for 9 days with and without the addition of GQDs. The longer time of the fibrillation is caused by the different aggregation feature of PSMα1 *in vitro* from PSMs in ECM of biofilm.^37^ Control peptides start forming short fibers in diameter of 11.4 ± 1.79 nm and in length of 134.4 ± 59.5 nm on Day 4, and the fibers were continuously elongated till Day 9 with similar diameter of 12.4 ± 0.82 nm (**Figure 3a, e**). The length of the elongated fibers on Day 9 was not measured as the fiber length exceed the field of view and was confounded by nonlinear geometry. Meanwhile, PSMα1 peptides were also incubated with GQDs at the concentrations of 50, 200 and 800 µg/mL. After incubation with GQDs for four days, small aggregates in diameter of 96.1 ± 49.98 nm, 127.9 ± 47.58 nm and 102.4 ± 48.90 nm instead of short fibers were found respectively in the peptide solution incubated with GQDs at the concentrations of 50, 200 and 800 µg/mL (**Figure 3b-d**). While incubated with 50 µg/mL GQDs after nine days, a few fibers in diameter of 10.39 ± 2.06 nm were formed as well (**Figure 3f**), including short fibers with length of 201.59 ± 64.4 nm and elongated fibers coiled together. At the same time, there are few short fibers and limited elongated fibers formed in diameter of 12.02 ± 2.79 nm in the sample of 200 µg/mL GQDs (**Figure 3g)**; however, there is only small aggregates in diameter of 32.28 ± 11.03 nm observed in the TEM images with 800 µg/mL GQDs (**Figure 3h**). In all the images, the contrast of the agglomerates was enhanced with increasing GQD concentration, which is associated with the tendency of GQDs to co-assemble with amyloid peptides. The delayed fibrillation for four days suggested that the kinetics of fibril formation changed in presence of GQDs which matches the expectations of the formation of ‘decoy’ complexes. Similarly to NPs,^42^ these supramolecular structures affect the secondary structure of peptides that serve as nucleation centers for fibrillation. Thus, binding of GQDs to PSM monomers or small prefibrillar oligomers is likely to hinder the formation of critical nuclei and elongation of the fibrils.

**Figure 3.**
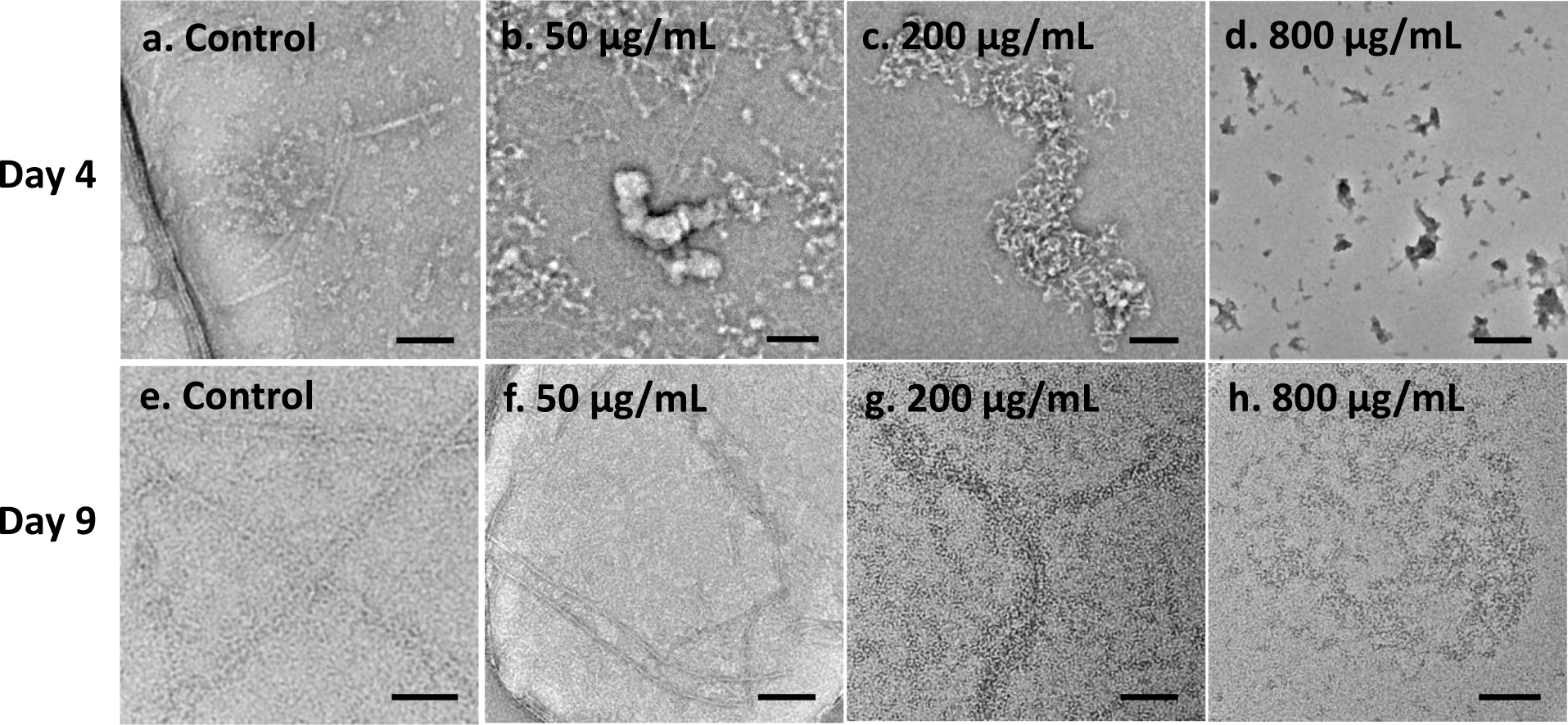
PSMα1 fibril formation with and without addition of GQDs. TEM images of PSMα1 peptides after addition of 0, 50, 200, 800 µg/mL GQDs **(a-d)** incubated for 4 days or **(e-h)** 9 days. Scale bars: 100 nm.

To verify this point, structural change in PSMα1 peptides interacting with GQDs were monitored for conformational conversion on Day 4 and Day 9 of incubation using CD spectrometry. As previously observed, there are two stages of conformational and morphological changes of peptide aggregation for PSM amyloid-like fibrils. Initially, the freshly dissolved PSMα1 peptides mainly contain *α-helix*es.^43^ The typical *β-sheet* signal with a negative band at ≈ 218 nm,^37^ appears after four days for PSMα1 without GQDs (**Figure 4a**). This suggested that formation of amyloid fibrils, which agreed with the result of ThT staining (**Figure S4**). Interestingly, secondary structures of PSMα1 incubated with GQDs showed enhancement of *β-turns* signal with a positive band at ≈208nm, in agreement with the theoretical peak of pure *β-turns* signal.^44,45^ Meanwhile, a broad peak from 250 nm and 375 nm was observed, corresponding to the absorption of GQDs onto peptide (**Figure S5**) and associated with the deformed (coiled) conformation of the graphene sheet.^15^ The combination of these two positive CD peaks of PSMα1 induced by GQDs in the solution are potentially contributed by the interaction between PSMα1 monomers and GQDs. Such interaction may change secondary structure of peptides incubated with GQDs, and/or enhance the CD signal of *turns* by GQD. Subsequently, the negative band at ≈ 218 nm of *β-sheet* was enhanced in PSMα1 self-assembling without GQDs after a nine-day incubation, which agrees with the significant increase in ThT fluorescence emission (**Figure 4b**, **S4**). The change of CD peak in PSMα1 control from Day 4 to Day 9 indicated the transition from lag phase of individual peptides and oligomer to fibril formation^42^. Fibrillation of PSMα1 incubated with GQDs was inhibited when the molar ratio of peptides and GQDs are equal or more than 1:1 (GQD concentration at 200 μg/mL). However, the fibrillation CD signal of PSMα1 was further interrupted when incubated with excess amount of GQDs (800 μg/mL) on both Day 4 and 9 (**Figure 4 a, b**).

**Figure 4.**
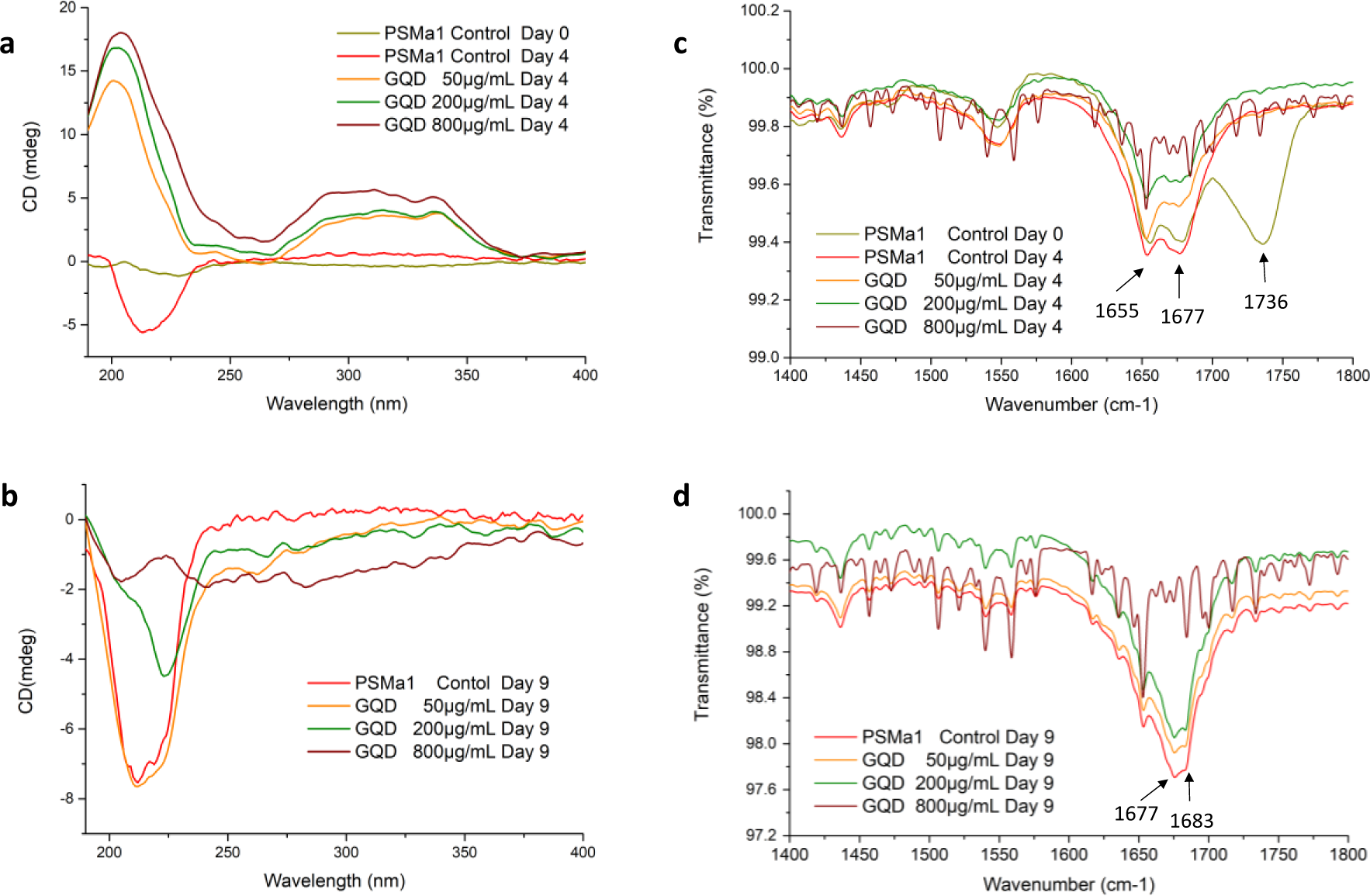
Secondary structure of synthetic PSMα1 peptides and fibrils with GQDs. **(a, b)** CD spectra of PSMα1 with/without addition of 50, 200, 800 µg/mL GQDs for **(a)** 4 days and **(b)** 9 days. FTIR spectra of PSMα1 with/without addition of 50, 200, 800 µg/mL GQDs for **(c)** 4 days and **(d)** 9 days. The intense signals between 1735–1740-cm^−1^, for control PSMα1 on Day 0, are indicative of C=O stretch of carboxylic group, which disappeared after incubation for four days in all samples. This is contributed by the minimum remains of trifluoroacetic acid (TFA) and hexafluoroisopropanol (HFIP), which cleave synthesized peptides to mono-dispersed peptide (**Figure S6**). This peak disappeared after 4 hours at room temperature and the remains did not change the pH of the peptide solution.

To confirm the change of secondary structure in protein, we monitored PSMα1 fibrillation with or without GQDs using attenuated total reflectance - Fourier transform infrared spectroscopy (ATR-FTIR) (**Figure 4c, d**). Deconvolution of the absorbance spectra in the amide I region^46^ is indicative of the individual secondary structure components and their relative contribution to the main signal. PSMα1 peptides incubated with or without addition of GQDs for four days display common bands both at ≈ 1655 cm^-1^ and ≈ 1677-cm^−1^. The peak at 1655 cm^-1^ corresponds to *α-helix* which appears between 1650 cm^-1^ and 1657cm^-1^ and overlapped with *turns* between 1655 cm^−1^ and 1675 cm^−1^. The latter is attributed to the weak peak of antiparallel β-sheet structure 1670-1690 cm^-1^. With addition of GQDs, antiparallel *β-sheet* were significantly decreased and indicated the inhibition of amyloid fibrillation at early stage of fibril growth. Meanwhile, the band at ≈ 1655 cm^−1^ right-shifted and became narrower, which verified the slight changes from *α-helix* to *β-turns* in PSMα1 dispersion. After 9 days, the *α-helix* peak decreased in all samples. At the same time, the peaks at 1670-1690 cm^-1^ increased in the samples of control, 50 and 200µg/mL of GQDs, indicating the formation of *β-sheet* nucleation centers for fibrillation. The peak intensity of *β-sheet* at 1677 cm^−1^ and 1683 cm^−1^ became lower with presence of higher concentration of GQDs, which verifies the inhibitory effect of GQD on fibrillation.

Molecular dynamics (MD) simulations were used to study the interactions between GQD and PSMα1 by monitoring the changes in secondary structure of the individual peptides. PSMα1 peptide contains 21 amino acids and consist of a single amphipathic *helix* with a slight bend near the *N*- and *C*-terminal (**Figure 5a**).^47^ Eight different simulations, with the GQD initially placed at the vertices of a cuboid encasing a single PSMα1, were tested to study the interactions between the two molecules. The results indicate that electrostatic interactions between the positive charged residues near the PSMα1 *N*-terminal, methionine (MET) and lysine (LYS), and –COO^-^ groups present on the edges of GQD lead to the formation of stable conformational complexes (**Figure 5b**). The simulations show a large pool of such complexes, where the position and the distance of the carboxylic groups affect the structure of the aggregate. Therefore, due to the relative high number of stable configurations and the slow transition among them, we cannot make conclusive remarks about the details of the most stable complexes. However, the analysis of the PSMα1 secondary structure changes due to the GQD-peptide interactions is revealing. The PSMα1-GQD complex, shows an increase of 311.0 % in the number of *turn* motifs compared to the isolated peptide (**Figure 5c**), which is consistent with the enhanced CD signal of *β-turns* at ≈208 nm. At the same time, the number of peptide configurations containing both *helix* and *coils* decreases by 3.0 %, but the distributions are affected differently: the residues involved in *helix* secondary structure decrease in both number and frequency, while the variance of the number of residues in a *coil* configuration increase (**Figure 5d, e**). In other words, the peptide losses α*-helix* structure in favor of the formation of *β-turns* or *coils*. Overall, the formation of stable PSMα1/GQD complexes by electrostatic interactions leads to relevant changes in the secondary structure and peptide motifs, and therefore we expect the presence of GQDs to affect unfavorably the organization of PSMα1 peptides on a larger scale. Specifically, the increase in structural disorder alters the initial stage of PSMα1 fibrillation thus inhibit the formation of nucleation centers.

**Figure 5.**
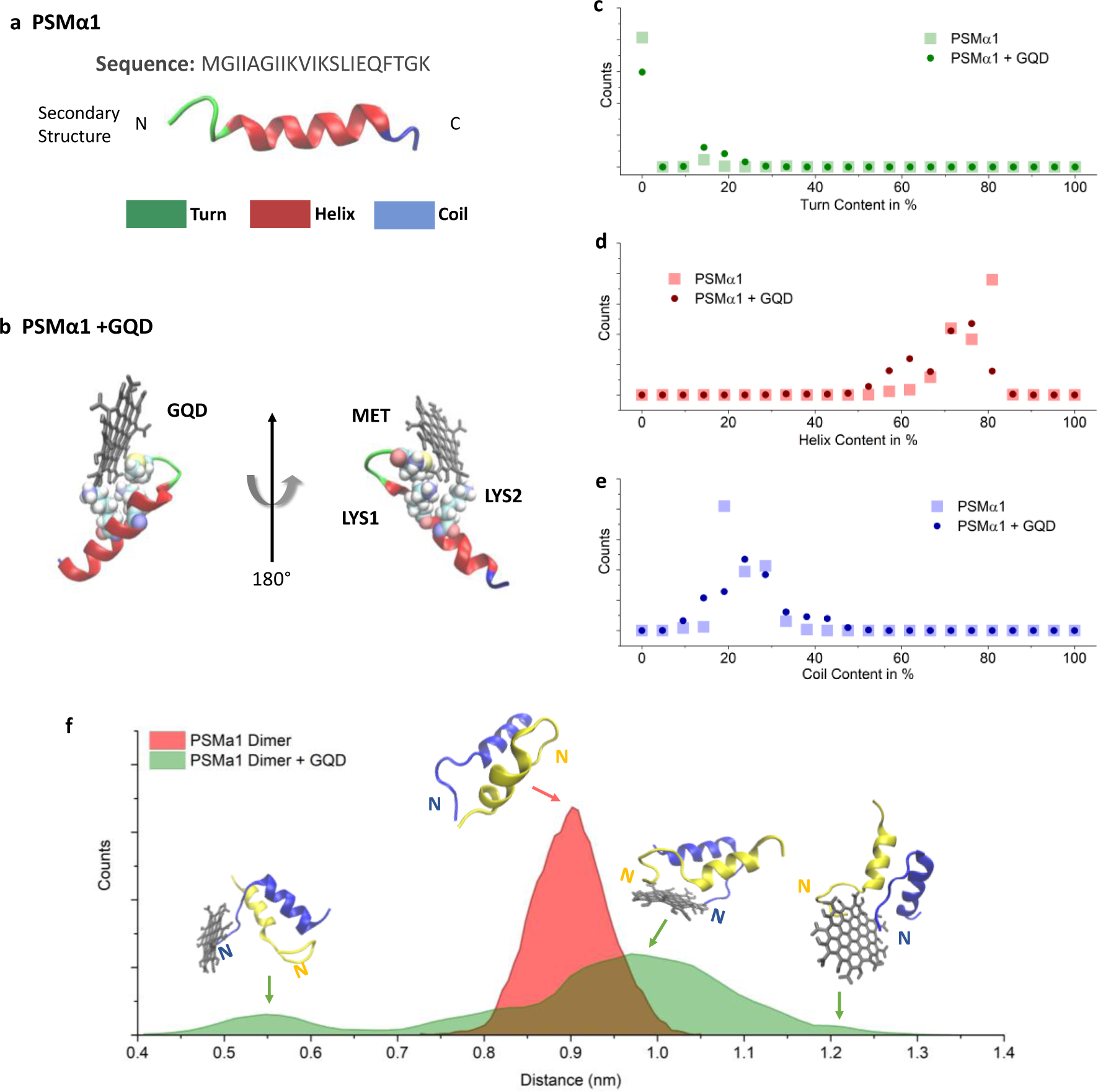
MD simulation of PSMα1 and GQDs. Schemes of (**a**) PSMα1^47^ and (**b**) PSMα1 and GQD complex. *β-turns* are shown in green, *alpha helix* in red and *random coils* in blue. **(c-e)** Histogram of secondary structure amounts in PSMα1 in the GQD/PSMα1 complex. (**f**) Histogram of the center of mass distance between two PSMα1 units showing the distance distribution with and without a GQD molecule. N: *N*-terminal of PSMα1; C: *C-*terminal of PSMα1.

MD simulations were also used to elucidate the impact of GQDs on the assembly of PSMα1 monomers. Specifically, the statistics of the inter-peptide distance (center of mass) were collected and analyzed (**Figure 5f**). The results show that the average distance increased from 0.76 ± 0.062 nm to 1.03 ± 0.055 nm when a GQD is in close proximity. Moreover, the distribution changes from a relatively symmetric monomodal bell-shaped curve centered at 0.9 nm to a distribution with three peaks (at 0.57, 0.97 and 1.2 nm) with a wide shoulder at 0.8 nm. While each peak is composed by a variety of configurations (representative complexes are shown in correspondence of each peak in **Figure 5f**), the analysis of the trajectories suggests that the inter-peptide *N*-terminal/*C*-terminal interaction, responsible for the dimer stability is disrupted by the presence of the GQD. Additionally, as the GQD interacts with *N*-terminal of a peptide, it alters the contact angle of the peptide backbones and causes the groups close to the *N*-terminal to stretch. Overall, the interaction between GQD and PSM peptides strongly suggests a frustration of the amyloid fibrillation, as observed experimentally (**Figure 6a**), caused by the GQDs docking near the *N*-terminus of the peptides and by changes in the secondary structure (**Figure 6b, c)**.

**Figure 6.**
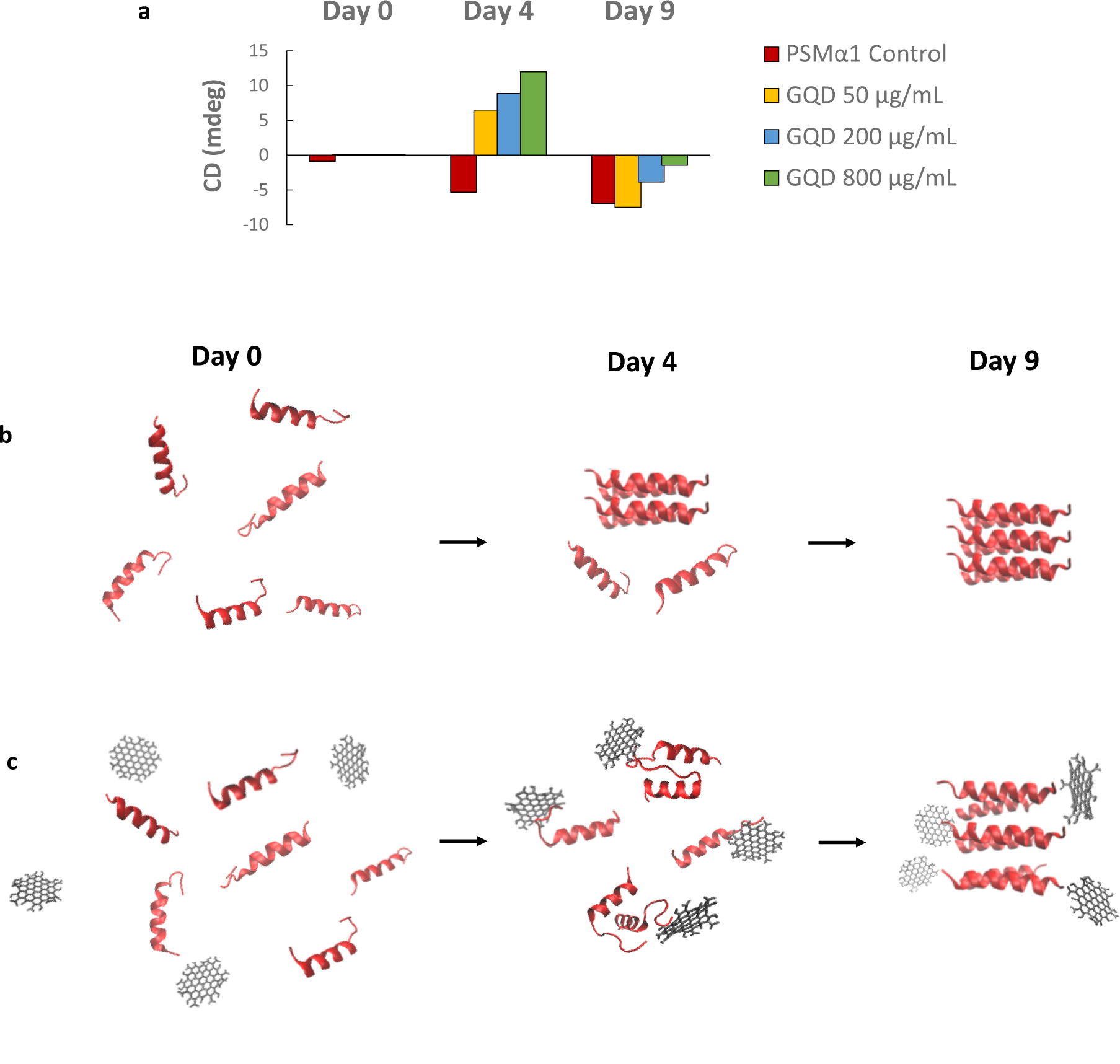
Schematics of the inhibiting effect of GQDs on PSM peptide fibrillation. **(a)** The timeline of CD intensity at the typical *β-sheet* signal, a negative band at ≈ 218 nm indicates the process of PSM peptide fibrillation with/without addition of 50, 200, 800 µg/mL GQDs. **(b, c)** GQDs interfere on *N-*terminal of PSM monomers or dimers and thereby inhibits fibrillation.

## CONCLUSIONS

Amyloid-rich biofilms are exceptionally resistant to chemical disruption^32,48^ and the ability to disperse such biofilms as a potential therapeutic alone or in combination with other antimicrobials is worthy of consideration. Here we identify GQDs as a potential therapeutic for this purpose. The specificity for amyloid-rich biofilms and lack of direct bacterial toxicity implicate amyloid-forming PSM peptides in the ECM as the molecular target for GQDs. This mechanism was confirmed through experiments with isolated biofilm ECM and synthetic PSMα1 as well as MD simulations. We conclude that GQDs disperse *S. aureus* biofilms by competitive assembly with amyloid peptides. The targeted disruption of functional amyloid formation likely results in increased free or oligomeric PSM peptides.^32^ The combination of reduced amyloid mediated biofilm stabilization and increased monomeric surfactant PSMs – which have been implicated in bacterial/ECM detachment^35^ – drives biofilm dispersion. In addition, such supramolecular complexes may also interfere with QS due to increased concentration of PSM monomers/oligomers that are tightly regulated by the Agr QS system.^48,49^ Considering the broader implications of this work, additional studies on the interaction between GQDs and amyloid-forming peptides in other species (e.g., *Escherichia coli* and *Pseudomonas spp*) will provide new approaches and insights into the potential manipulation of microbial communities. Further engineering of GQDs for enhanced association with amyloid peptides may lead to diverse effects on a wide range of biological processes, biomolecular functions, and signaling pathways.

## METHODS

### Synthesis and Characterization of GQDs

GQDs were synthesized by a modified protocol through a top-down “oxidation-cutting” process.^15^ 50 mg of carbon fibers was dispersed into a 4 mL mixture of sulfuric acid and nitric acid (3:1 v/v). The prepared solution was sonicated for 2 h and mechanically stirred for 24 h at 80 °C. Afterward, the mixture was cooled and diluted with deionized (DI) water to the concentration of 0.15 mg/mL. Sodium hydroxide was added into the solution to adjust the pH value to neutral. The mixture was further dialyzed in DI water for 3 days to obtain the final product. The size and shape of GQDs were characterized by TEM (JEOL 3011 HREM). The element analysis was performed by XPS (Kratos, Axis Ultra XPS). The light emission properties of GQDs and their assemblies were investigated by fluorescence spectroscopy (Horiba, Fluoromax-3).

### Biofilm Culture

*S. aureus* USA300 from −80°C glycerol stock was plated on tryptic soy agar. Single colony inoculates were grown to mid-log, optical density 0.4 - 0.8 at 600 nm (OD_600_). Mid-log cultures were diluted to OD_600_ = 0.02 for seeding biofilms. For treating intact biofilms with GQDs 100 µl of this suspension had 10^6^ *S. aureus* cells which were seeded into Stovall triple chamber flow cells. Biofilms were allowed to grow for 72 hours with either TSBG or PNG media flowing at 13 μl/min. Then 0, 50, or 500 µg/mL of GQDs were added and the flow was resumed for an additional 24 h. For isolating biofilm ECM, 100 µl suspensions containing 10^6^ *S. aureus* were seeded on to polystyrene culture plates. Biofilms were then grown under static conditions for 5 days. All bacterial cultures were maintained at 37 °C.

### Peptide Preparation

PSMα1 was purchased from ChinaPeptides (Shanghai, China) with a purity >90%. They were dissolved to a final concentration of 0.5 mg/mL in a 1:1 mixture of trifluoroacetic acid (TFA) and hexafluoroisopropanol (HFIP). Peptides were then sonicated for 10 minutes and incubated for 1 h at room temperature. Stock solutions were divided into aliquots, solvent TFA/HFIP dried with a SpeedVac (Thermo Scientific, USA) at room temperature and stored at −80 °C. Samples were suspended in anhydrous dimethyl sulfoxide (5%) and sonicated for 10 minutes. This preparation yielded PSMs in monomeric form.

### Thioflavin T (ThT) Assay

Purified PSM solution was mixed with a ThT stock solution (prepared in 50 mM NaCl, 20 mM Tris, pH 8) to a final concentration of 2–4 μM peptides, 0.1-0.25 mM ThT. The final volume was 200 μl, and measurements were performed in 96-well plates with a clear bottom. Fixed excitation/emission filter of 430/480 nm (Ex/Em), respectively. Data was collected every hour for 15 hours.

### Circular dichroism (CD) Spectrometry

CD spectra were measured in a Jasco-815 spectropolarimeter (Jasco, Japan) thermostated at 25 °C. Aggregated peptides were prepared at 20 μM and measured immediately. Spectra were recorded from 190 to 400 nm, at 0.5 nm intervals, 1 nm bandwidth, and a scan speed of 100 nm/min.

### Confocal Imaging

For biofilm visualization by confocal laser scanning microscopy (Nikon A1 plus), stains were added to TSBG or PNG within the flow cells at the conclusion of the biofilm culture experiment. The stains were incubated under static conditions in the flow cells for 20 minutes. Wheat Germ Agglutinin (WGA) was used to visualize PIA while cells were stained with DAPI (4′,6-diamidino-2-phenylindole). Three sets of samples were prepared at different time periods. Nine to fifteen images were taken in each sample for image analysis.

### Image Analysis

Digital images captured from confocal fluorescence microscopy and SEM were processed in ImageJ in the following manner: (1) Using process subtract background, background corrections were made. (2) Using image adjust threshold, a constant threshold level was set across a set of conditions to ensure that enough area of signal was present to identify the target including PIA, bacterial cells. (3) A constant area was applied to measure the area and integrated fluorescent density of the measured content.

### Biofilm ECM Isolation

Biofilm ECM was harvested by disrupting biofilms using vortex and sonication as previously described.^32^ Briefly fibers were collected after 5-day static growth by scraping biofilms into 3 mL of potassium phosphate buffer (50 mM, pH 7.4). The biofilm suspensions were homogenized using a tissue homogenizer (Power 4, 3 minutes) to shear fibers free from the cell walls. Supernatants were clarified by repeated (×2) centrifugation at 4,000 g for 15 minutes to remove cells. The cell-free supernatant was incubated in 200 mM NaCl for 24 hours at room temperature. The fibers were isolated using Millipore Amicon Ultra Centrifugal Filter Units with a pore size of 30 kDa.

### Scanning Electron Microscopy (SEM)

Biofilms were cultured on sample holders as the standard procedure. GQDs (500 µg/mL) were added incubated for 24 hours at 37 °C. Similarly, isolated ECM was placed on sample holders and treated with GQDs. All samples were fixed in glutaraldehyde, serially dehydrated in ethanol, air dried at room temp, sputter-coated with gold and visualized using AMRAY 1910 Field Emission Scanning Electron Microscope.

### Transmission Electron Microscopy (TEM)

The TEM employed in the characterization is JEOL 3011 HREM. Carbon-coated 400 mesh copper TEM grids were placed coated-side-down for 60 s onto sample drops (10 µl) of PSMα1 solution with/without addition of 50, 200, 800 µg/mL GQDs for 4 and 9 days. The grids were then retrieved, washed with deionized water (two droplets). The sample was stained with 1% (w/v) uranyl acetate for 40 s. Grids were then blotted and air-dried before imaging. For imaging cells with amyloid containing ECM, biofilms were scrapped directly on to copper TEM grids, stained with uranyl acetate, dried and imaged.

### Molecular Dynamics (MD) Simulation

All the simulations were performed with the NAMD in explicit water using the classical all-atom force field CHARMM. A time step of 1 fs was employed to integrate the equations of motion. A cutoff of 1.2 nm was used in conjunction with the Particle Mesh Ewald method to evaluate long-range columbic forces. The systems were minimized for 1000 steps before the production runs. Single peptide simulations for a total or more than 650 ns, were performed in a canonical ensemble (300 K, with a time constant of 1 ps) in a cubic simulation box with side 7.5 nm. Production runs of the PSMα1 dimers were performed in a canonical ensemble in a cubic simulation box with a side of 10 nm.

## ACKNOWLEDGEMENTS

This project was sponsored by the University of Michigan Mcubed Award and Defense Advanced Research Projects Agency (DARPA) HR00111720067 “Electromagnetic Processes and Normal Modes in Bacterial Biofilms”. Parts of the work were also supported by NIH under grant K08 AI128006 and NSF project “Energy- and Cost-Efficient Manufacturing Employing Nanoparticles” NSF 1463474. We are acknowledging the financial and programmatic support from the University of Michigan College of Engineering’s Blue Sky Initiative. We are also acknowledging the support from UM electron microscopy facility MC2 for its assistance with electron microscopy, and for the NSF grant #DMR-9871177 for funding of the JEOL 2010F analytical electron microscope used in this work.

## AUTHOR CONTRIBUTIONS

Y.W., N.A.K., A.V. and J.S.V originated and conceptualized the study. Y.W. and U.K. designed the experiments. Y.W. carried out the experiments and the MD simulation. U.K. prepared the biofilms and isolated ECM. Z.Q. synthesized and characterized the GQDs. P.E. contributed to design and analysis of the MD simulation. C.A. contributed to preparation of the PNG biofilms. N.A.K., A.V. and J.S.V supervised all the work. Y.W. prepared the manuscript and all authors contributed to data interpretation, discussions and writing.

## REFERENCES

(1) Hill, E. C.; Hill, G. C. Microbial Contamination and Associated Corrosion in Fuels, during Storage, Distribution and Use. Adv. Mater. Res. 2008, 38, 257–268.

(2) Porcelli, N.; Judd, S. Chemical Cleaning of Potable Water Membranes: A Review. Sep. Purif. Technol. 2010, 71, 137–143.

(3) Costerton, J. W. Bacterial Bofilms: A Common Cause of Persistent Infections. Science (80-.). 1999, 284, 1318–1322.

(4) Castaño-Rodríguez, N.; Goh, K.-L.; Fock, K. M.; Mitchell, H. M.; Kaakoush, N. O. Dysbiosis of the Microbiome in Gastric Carcinogenesis. Sci. Rep. 2017, 7, 15957.

(5) Davey, M. E.; O’toole, G. A. Microbial Biofilms: From Ecology to Molecular Genetics. Microbiol. Mol. Biol. Rev. 2000, 64, 847–867.

(6) Neethirajan, S.; Clond, M. A.; Vogt, A. Medical Biofilms - Nanotechnology Approaches. J. Biomed. Nanotechnol. 2014, 10, 2806–2827.

(7) Hall-Stoodley, L.; Costerton, J. W.; Stoodley, P. Bacterial Biofilms: From the Natural Environment to Infectious Diseases. Nat. Rev. Microbiol. 2004, 2, 95–108.

(8) Branda, S. S.; Vik, Å.; Friedman, L.; Kolter, R. Biofilms: The Matrix Revisited. Trends Microbiol. 2005, 13, 20–26.

(9) Tursi, S. A.; Tükel, Ç. Curli-Containing Enteric Biofilms Inside and Out: Matrix Composition, Immune Recognition, and Disease Implications. Microbiol. Mol. Biol. Rev. 2018, 82.

(10) Olsen, I. Biofilm-Specific Antibiotic Tolerance and Resistance. Eur. J. Clin. Microbiol. Infect. Dis. 2015, 34, 877–886.

(11) Arciola, C. R.; Campoccia, D.; Speziale, P.; Montanaro, L.; Costerton, J. W. Biofilm Formation in Staphylococcus Implant Infections. A Review of Molecular Mechanisms and Implications for Biofilm-Resistant Materials. Biomaterials 2012, 33, 5967–5982.

(12) Brogden, K. A. Antimicrobial Peptides: Pore Formers or Metabolic Inhibitors in Bacteria. Nat. Rev. Microbiol. 2005, 3, 238–250.

(13) Wu, H.; Moser, C.; Wang, H.-Z.; Høiby, N.; Song, Z.-J. Strategies for Combating Bacterial Biofilm Infections. Int. J. Oral Sci. 2015, 7, 1–7.

(14) Romero, D.; Sanabria-Valentín, E.; Vlamakis, H.; Kolter, R. Biofilm Inhibitors That Target Amyloid Proteins. Chem. Biol. 2013, 20, 102–110.

(15) Suzuki, N.; Wang, Y.; Elvati, P.; Qu, Z.-B.; Kim, K.; Jiang, S.; Baumeister, E.; Lee, J.; Yeom, B.; Bahng, J. H.; et al. Chiral Graphene Quantum Dots. ACS Nano 2016, 10, 1744–1755.

(16) Sun, H.; Gao, N.; Dong, K.; Ren, J.; Qu, X. Graphene Quantum Dots-Band-Aids Used for Wound Disinfection. ACS Nano 2014, 8, 6202–6210.

(17) Zeng, Z.; Yu, D.; He, Z.; Liu, J.; Xiao, F.-X.; Zhang, Y.; Wang, R.; Bhattacharyya, D.; Tan, T. T. Y. Graphene Oxide Quantum Dots Covalently Functionalized PVDF Membrane with Significantly-Enhanced Bactericidal and Antibiofouling Performances. Sci. Rep. 2016, 6, 20142.

(18) Hui, L.; Huang, J.; Chen, G.; Zhu, Y.; Yang, L. Antibacterial Property of Graphene Quantum Dots (Both Source Material and Bacterial Shape Matter). ACS Appl. Mater. Interfaces 2016, 8, 20–25.

(19) Tang, Z.; Kotov, N. A.; Giersig, M. Spontaneous Organization of Single CdTe Nanoparticles into Luminescent Nanowires. Science (80-.). 2002, 297, 237–240.

(20) Zhou, Y.; Marson, R. L.; van Anders, G.; Zhu, J.; Ma, G.; Ercius, P.; Sun, K.; Yeom, B.; Glotzer, S. C.; Kotov, N. A. Biomimetic Hierarchical Assembly of Helical Supraparticles from Chiral Nanoparticles. ACS Nano 2016, 10.1021/ac.

(21) Park, J. Il; Nguyen, T. D.; de Queirós Silveira, G.; Bahng, J. H.; Srivastava, S.; Zhao, G.; Sun, K.; Zhang, P.; Glotzer, S. C.; Kotov, N. A. Terminal Supraparticle Assemblies from Similarly Charged Protein Molecules and Nanoparticles. Nat. Commun. 2014, 5.

(22) Huang, R.; Carney, R. P.; Stellacci, F.; Lau, B. L. T. Protein-Nanoparticle Interactions: The Effects of Surface Compositional and Structural Heterogeneity Are Scale Dependent. Nanoscale 2013, 5, 6928–6935.

(23) Lundqvist, M.; Stigler, J.; Elia, G.; Lynch, I.; Cedervall, T.; Dawson, K. A. Nanoparticle Size and Surface Properties Determine the Protein Corona with Possible Implications for Biological Impacts. Proc. Natl. Acad. Sci. U. S. A. 2008, 105, 14265–14270.

(24) Harkness, K. M.; Turner, B. N.; Agrawal, A. C.; Zhang, Y.; McLean, J. A.; Cliffel, D. E.; Haupt, K.; Mosbach, K.; Shin, H.; Jo, S.; et al. Biomimetic Monolayer-Protected Gold Nanoparticles for Immunorecognition. Nanoscale 2012, 4, 3843.

(25) Sun, J.; DuFort, C.; Daniel, M.-C.; Murali, A.; Chen, C.; Gopinath, K.; Stein, B.; De, M.; Rotello, V. M.; Holzenburg, A.; et al. Core-Controlled Polymorphism in Virus-like Particles. Proc. Natl. Acad. Sci. U. S. A. 2007, 104, 1354–1359.

(26) Yoo, S. Il; Yang, M.; Brender, J. R.; Subramanian, V.; Sun, K.; Joo, N. E.; Jeong, S.-H.; Ramamoorthy, A.; Kotov, N. A. Inhibition of Amyloid Peptide Fibrillation by Inorganic Nanoparticles: Functional Similarities with Proteins. Angew. Chemie Int. Ed. 2011, 50, 5110–5115.

(27) Carrillo-Carrion, C.; Atabakhshi-Kashi, M.; Carril, M.; Khajeh, K.; Parak, W. J. Taking Advantage of Hydrophobic Fluorine Interactions for Self-Assembled Quantum Dots as a Delivery Platform for Enzymes. Angew. Chemie Int. Ed. 2018, 57, 5033–5036.

(28) Mahmoudi, M.; Akhavan, O.; Ghavami, M.; Rezaee, F.; Ghiasi, S. M. A. Graphene Oxide Strongly Inhibits Amyloid Beta Fibrillation. Nanoscale 2012, 4, 7322.

(29) Liu, Y.; Xu, L.-P.; Dai, W.; Dong, H.; Wen, Y.; Zhang, X. Graphene Quantum Dots for the Inhibition of β Amyloid Aggregation. Nanoscale 2015, 7, 19060–19065.

(30) Wang, J.; Cao, Y.; Li, Q.; Liu, L.; Dong, M. Size Effect of Graphene Oxide on Modulating Amyloid Peptide Assembly. Chem. - A Eur. J. 2015, 21, 9632–9637.

(31) Kim, D.; Yoo, J. M.; Hwang, H.; Lee, J.; Lee, S. H.; Yun, S. P.; Park, M. J.; Lee, M.; Choi, S.; Kwon, S. H.; et al. Graphene Quantum Dots Prevent α-Synucleinopathy in Parkinson’s Disease. Nat. Nanotechnol. 2018, 13, 812–818.

(32) Schwartz, K.; Syed, A. K.; Stephenson, R. E.; Rickard, A. H.; Boles, B. R. Functional Amyloids Composed of Phenol Soluble Modulins Stabilize Staphylococcus Aureus Biofilms. PLoS Pathog. 2012, 8, e1002744.

(33) Periasamy, S.; Chatterjee, S. S.; Cheung, G. Y. C.; Otto, M. Phenol-Soluble Modulins in Staphylococci. Commun. Integr. Biol. 2012, 5, 275–277.

(34) Bleem, A.; Francisco, R.; Bryers, J. D.; Daggett, V. Designed α-Sheet Peptides Suppress Amyloid Formation in Staphylococcus Aureus Biofilms. npj Biofilms Microbiomes 2017, 3, 16.

(35) Cheung, G. Y. C.; Joo, H.-S.; Chatterjee, S. S.; Otto, M.; Fd, L.; Fd, L.; Rp, N.; M, O.; M, O.; M, O.; et al. Phenol-Soluble Modulins – Critical Determinants of Staphylococcal Virulence. FEMS Microbiol. Rev. 2014, 38, 698–719.

(36) Tayeb-Fligelman, E.; Tabachnikov, O.; Moshe, A.; Goldshmidt-Tran, O.; Sawaya, M. R.; Coquelle, N.; Colletier, J.-P.; Landau, M. The Cytotoxic Staphylococcus Aureus PSMα3 Reveals a Cross-α Amyloid-like Fibril. Science 2017, 355, 831–833.

(37) Marinelli, P.; Pallares, I.; Navarro, S.; Ventura, S.; Rubin, R. J.; Gordon, R. J.; Lowy, F. D.; Boucher, H.; Miller, L. G.; Razonable, R. R.; et al. Dissecting the Contribution of Staphylococcus Aureus α-Phenol-Soluble Modulins to Biofilm Amyloid Structure. Sci. Rep. 2016, 6, 34552.

(38) Pollitt, E. J. G.; Crusz, S. A.; Diggle, S. P.; Josenhans, C.; Suerbaum, S.; O’Toole, G. A.; Kolter, R.; Henrichsen, J.; Kaito, C.; Sekimizu, K.; et al. Staphylococcus Aureus Forms Spreading Dendrites That Have Characteristics of Active Motility. Sci. Rep. 2015, 5, 17698.

(39) Pauling, L.; Corey, R. B.; Xi, W.; Luo, F.; Zhang, X.; Zou, M.; Cao, M.; Hu, J.; Wang, W.; Wei, G.; et al. Configurations of Polypeptide Chains With Favored Orientations Around Single Bonds: Two New Pleated Sheets. Proc. Natl. Acad. Sci. U. S. A. 1951, 37, 729–740.

(40) Elvati, P.; Baumeister, E.; Violi, A. Graphene Quantum Dots: Effect of Size, Composition and Curvature on Their Assembly. RSC Adv. 2017, 7, 17704–17710.

(41) Das, T.; Kutty, S. K.; Tavallaie, R.; Ibugo, A. I.; Panchompoo, J.; Sehar, S.; Aldous, L.; Yeung, A. W. S.; Thomas, S. R.; Kumar, N.; et al. Phenazine Virulence Factor Binding to Extracellular DNA Is Important for Pseudomonas Aeruginosa Biofilm Formation. Sci. Rep. 2015, 5, 8398.

(42) Arosio, P.; Knowles, T. P. J.; Linse, S. On the Lag Phase in Amyloid Fibril Formation. Phys. Chem. Chem. Phys. 2015, 17, 7606–7618.

(43) Sreerama, N.; Woody, R. W. Estimation of Protein Secondary Structure from Circular Dichroism Spectra: Comparison of CONTIN, SELCON, and CDSSTR Methods with an Expanded Reference Set. Anal. Biochem. 2000, 287, 252–260.

(44) Bush, C. A.; Sarkar, S. K.; Kopple, K. D. Circular Dichroism of β Turns in Peptides and Proteins. Biochemistry 1978, 17, 4951–4954.

(45) Greenfield, N. J. Using Circular Dichroism Spectra to Estimate Protein Secondary Structure. Nat. Protoc. 2007, 1, 2876–2890.

(46) Spectroscopic Methods for Analysis of Protein Secondary Structure. Anal. Biochem. 2000, 277, 167–176.

(47) Towle, K. M.; Lohans, C. T.; Miskolzie, M.; Acedo, J. Z.; van Belkum, M. J.; Vederas, J. C. Solution Structures of Phenol-Soluble Modulins Α1, Α3, and Β2, Virulence Factors from Staphylococcus Aureus. Biochemistry 2016, 55, 4798–4806.

(48) Rice, K. C.; Murphy, E.; Yang, S.-J.; Bayles, K. W.; Smeltzer, M. S.; Projan, S. J.; Beenken, K. E.; Cassat, J.; Palm, K. J.; Dunman, P. M. Transcriptional Profiling of a Staphylococcus Aureus Clinical Isolate and Its Isogenic Agr and SarA Mutants Reveals Global Differences in Comparison to the Laboratory Strain RN6390. Microbiology 2006, 152, 3075–3090.

(49) Xu, T.; Wang, X.-Y.; Cui, P.; Zhang, Y.-M.; Zhang, W.-H.; Zhang, Y. The Agr Quorum Sensing System Represses Persister Formation through Regulation of Phenol Soluble Modulins in Staphylococcus Aureus. Front. Microbiol. 2017, 8, 2189.

