## Supplemental Information for "Anti-biofilm Activity of Graphene Quantum Dots via Self-Assembly with Bacterial Amyloid Proteins"

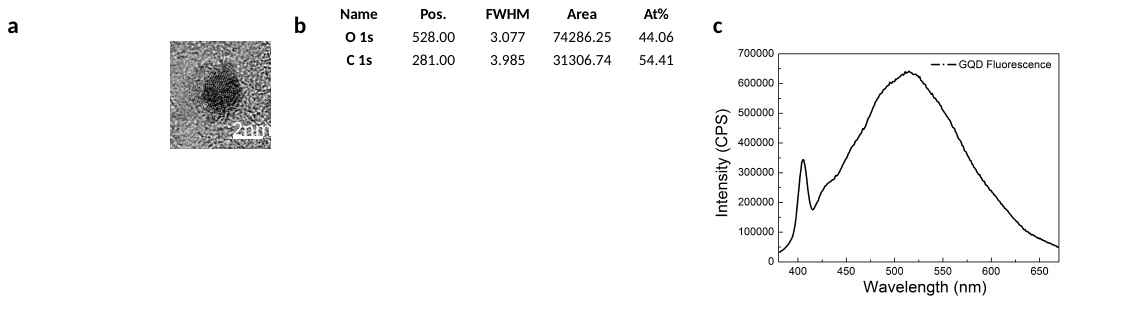

Figure S1 Characterization of as synthesized GQDs. (a) TEM images of GQDs as synthesized (b) X-ray photoelectron spectroscopy (XPS) of GQDs (c) Fluorescence of GQD exited by 350 nm photons.

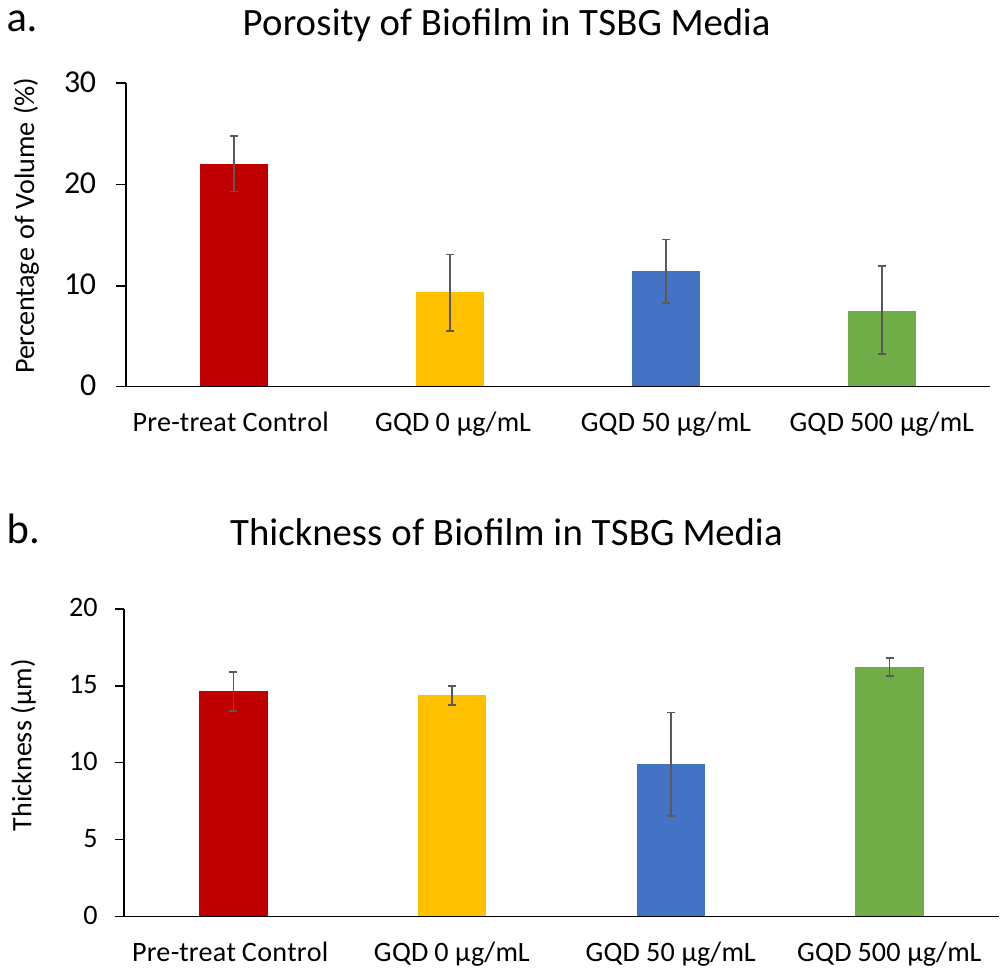

Figure S2 Quantitative image analysis derived from confocal microscopy of TSBG grown *S. aureus* biofilms. (a) Porosity of biofilm grown in TSBG medium as control as well as treated with 0 µg/mL, 50 µg/mL, and 500 µg/mL GQDs. (b) Thickness of TSBG grown biofilm as control as well as treated with 0 µg/mL, 50 µg/mL, and 500 µg/mL GQDs. Pre-treat control group represent the biofilm at the time before adding GQDs.

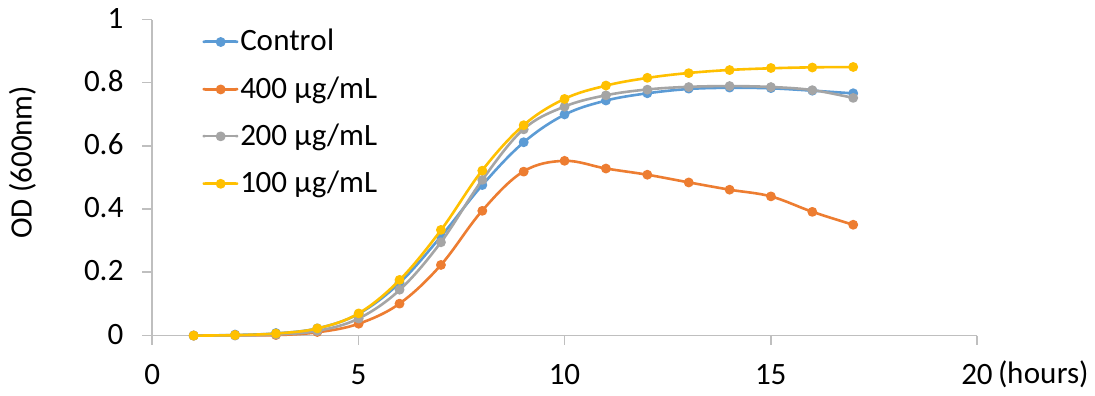

Figure S3 Planktonic growth curve for *S. aureus* in the presence of increasing concentrations of GQDs.

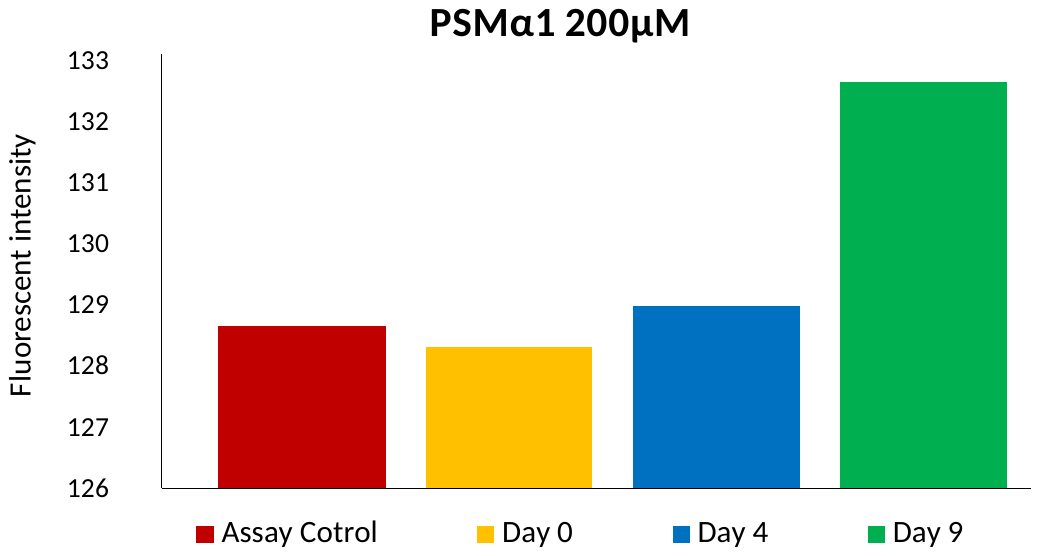

Figure S4 ThT staining of PSMα1 after incubation for 0, 4 and 9 days.

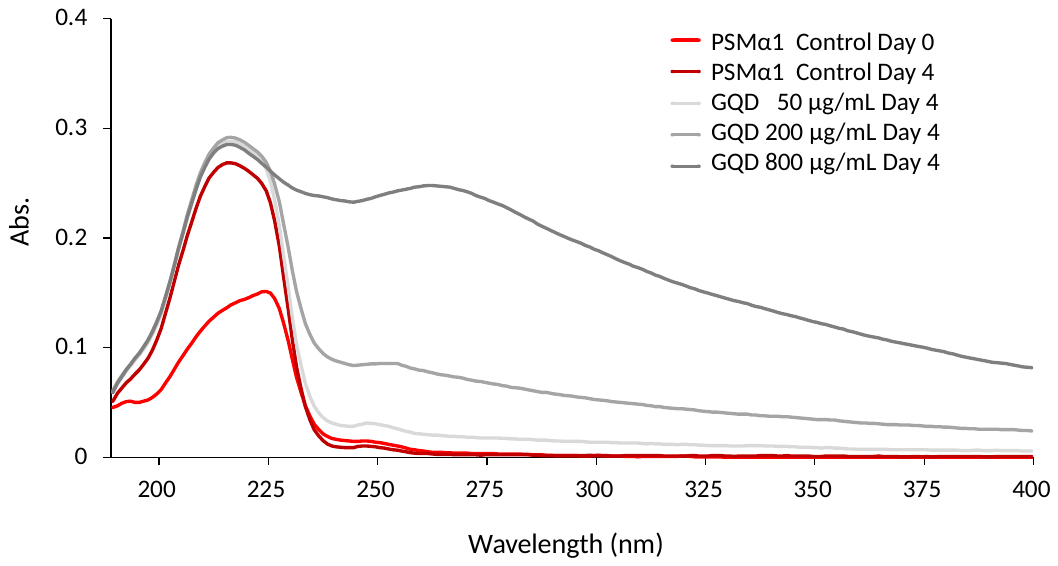

Figure S5 Absorption spectra of 200 µM PSMα1 with or without GQDs (50, 200, 800 µg/mL) for 0 and 4 days. The measured solution was diluted by 10 times of original incubated solution.

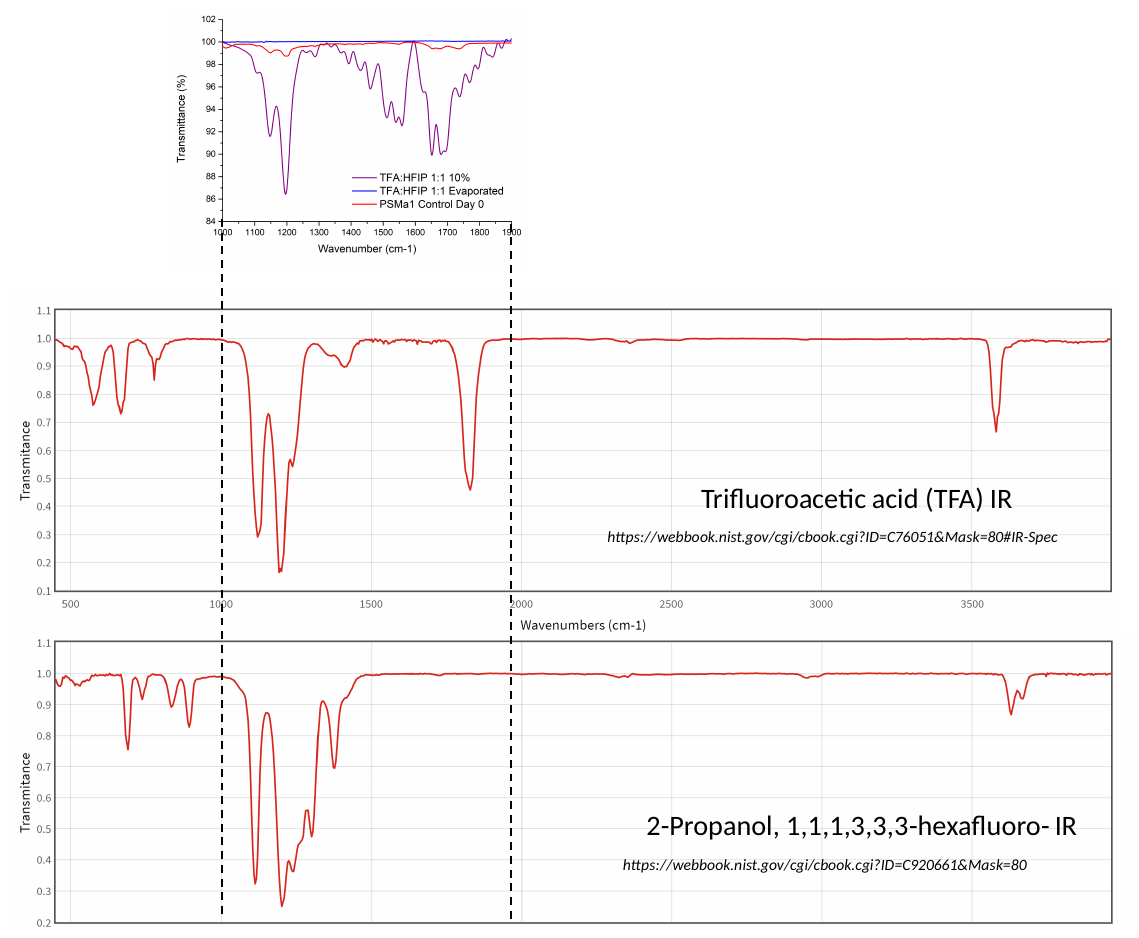

**Figure S6** FTIR of TFA and HFIP control solution comparing to their reference spectra and the spectrum of PSMα1 control sample on Day 0.
